# Predicting patient-specific radiotherapy responses in head and neck cancer to personalize radiation dose fractionation

**DOI:** 10.1101/630806

**Authors:** Heiko Enderling, Enakshi Sunassee, Jimmy J. Caudell

## Abstract

Human papillomavirus (HPV) related oropharyngeal cancer (OPC) is one of the few types of cancers increasing in incidence. HPV+ OPC treatment with radiotherapy (RT) provides 75-95% five-year locoregional control (LRC). Why some but not all patients with similar clinical stage and molecular profile are controlled remains unknown. We propose the proliferation saturation index, PSI, as a mathematical modeling biomarker of tumor growth and RT response. The model predicts that patients with PSI<0.75 are likely to be cured by radiation, and that hyperfractionated radiation could improve response rates for patients with higher PSI that are predicted to fail standard of care RT. Prospective evaluation is currently ongoing.

## I. INTRODUCTION

Radiotherapy (RT) is one of the single most commonly delivered oncologic treatments, utilized in over half of all cancer patients at some point in their care. In head and neck cancer, over 80% of patients receive RT, either as curative therapy alone or in combination with surgery and/or chemotherapy [1]. Human papillomavirus related (HPV+) oropharyngeal cancer (OPC) are one of the few types of cancers increasing in incidence. Treatment of OPC patients almost always includes radiotherapy (RT) either as curative therapy alone or in combination with surgery and/or chemotherapy. Standard RT delivers 2 Gy daily fractions for a total of 66-70 Gy (6-7 weeks), providing five-year locoregional control (LRC) for 75-95% of HPV+ OPC patients. Total RT dose and dose fractionation are based on clinical outcome data from large clinical trials testing intensification methods prior to the HPV era, resulting in a “one size fits all” approach. In current radiation oncology practice, there exists no explanation for why two patients with similar histology, primary site, and clinical stage would have different responses and outcomes. Reliable biomarkers and frameworks are needed to predict responses to personalized dose and fractionation of RT based on individual tumor features. Despite a long history of medical physics and physical concepts centered around radiation dose delivery technology and safety, few inroads have been made to synergize biological and quantitative approaches with radiation biology and radiation oncology methodologies to optimize RT and treatment personalization.

Precision medicine tools such as genomics, radiomics, and mathematical modeling could help personalize and adapt RT for each patient to improve cancer outcomes [2]. Prior work by our group suggests that a robust mid-treatment nodal response is prognostic for ultimate outcome in OPC patients. Patients with a nodal volume regression of > 32% at week 4 (20 fractions of RT) have 100% disease-free survival at 4-years follow up, compared with 77.3% (p = 0.02) for those with regression ≤ 32% (**Fig. 1A**,**B**)[3]. To prospectively predict RT response on a per-patient basis, our group has pioneered the development of the non-invasive imaging-derived proliferation saturation index (PSI) that can be calculated from routinely collected radiological images prior to therapy at diagnosis and RT planning session [4-5]. PSI provides an estimation of the tumor microenvironment-modulated radiosensitive proliferating subpopulation in solid tumors, and could predict RT responses and identify individual patient candidates for alternate RT fractionation protocols.

**Figure 1.**
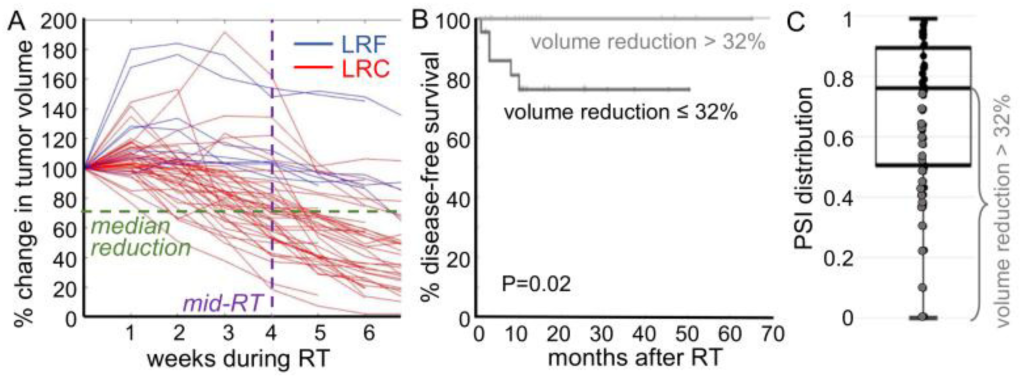
**A.** Change in volume during RT for N=49 patients. Median volume reduction by mid-RT (week 4) is 32%. Red trajectories: locoregionally controlled (LRC) patients. Blue: locoregional failure (LRF). **B.** Disease free survival by mid-RT volume reduction. **C.** Distribution of PSI for all patients.

## II. METHODS

Based on established logistic tumor growth dynamics concepts, we introduced a patient-specific proliferation saturation index (PSI) [4] to derive the fraction of radiation-sensitive proliferating population of cancer cells within an imaging-derived tumor volume prior to treatment:

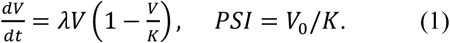

PSI is denoting the tumor volume prior to therapy (*V*_*0*_) to carrying capacity ratio. The carrying capacity denotes the tumor-extrinsic tissue-environmental properties of the patient that influence tumor growth dynamics, including the established oxygen and nutrients supply through tissue vascularization, removal of metabolic waste products, and evasion of immune surveillance.

Radiation response is modeled as decrease in tumor volume such that the volume after irradiation (V_postIR_) is the volume prior to radiation reduced by cell death of proliferating cells at rate:

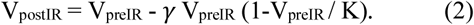

From radiation-induced volume reduction rate *γ* we can derive the OPC specific linear-quadratic radiation sensitivity parameters α and β with *γ = 1 -exp(-αd-βd*^*2*^), where *d* is the radiation dose. In this model, tumor growth rate (or volume doubling time) and radiation sensitivity are assumed uniform across patients, and tumor carrying capacity is modeled as the only patient-specific parameter that yields a patient-specific pre-treatment PSI. It follows from that larger PSI reflect a low proliferating cell fraction and, thus, potentially treatment refractory tumors, whereas tumors with low PSI are more proliferative and potentially radiosensitive. As such, two patients that present with similar tumor volume could have a different PSI, which results in different responses to the same RT protocol (**Fig. 2A**).

**Figure 2:**
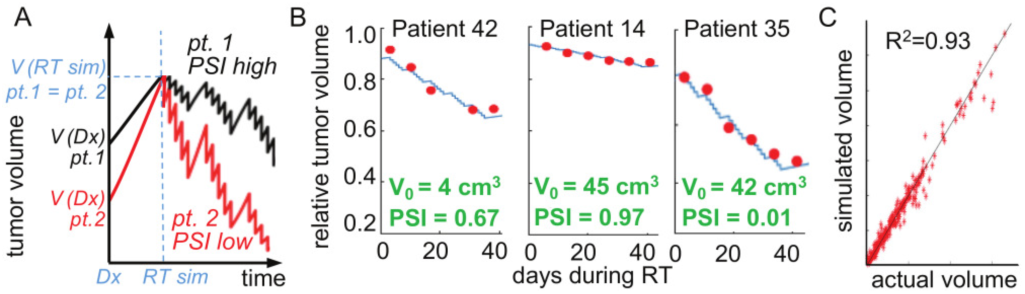
**A.** Conceptual PSI framework. Two patients with identical tumor volume at time of RT simulation (RT sim) can have different PSI due to different radiographic volumes at diagnosis. Higher PSI value predicts lower tumor volume reduction during RT. **B.** Representative examples of PSI model fit to clinical data. Red circles: radiographic volumes, blue curves: model fit. Patient-specific initial tumor volume (V_0_) and PSI shown in green. **C.** High correlation of simulated vs. actual weekly measured tumor volumes of N=49 OPC patients during RT (R^2^=0.93).

## III. RESULTS

PSI as the sole patient-specific marker of RT response was able to fit the data of N=49 OPC patients during RT with high accuracy (R^2^=0.93, **Fig. 2B, C**) with λ=0.02 day^-1^and *γ*=0.14. The distribution of patient-specific PSI is shown in **Fig. 1C**. Patients with >32% tumor volume reduction by week 4 had PSI<0.75.

RT with standard of care 2 Gy daily yielded a mid-treatment tumor volume reduction >32% in 24 of 49 patients (49%). Simulations of the PSI model suggest that 31 of 49 patients (63.3%) would achieved tumor volume reductions > 32% by week four with 1.2 Gy twice daily (B.I.D., **Fig. 3**). Patients with PSI = (0.75,0.85) are most likely to achieve the 32% mid-treatment response with hyperfractionation compared to standard of care.

**Figure 3:**
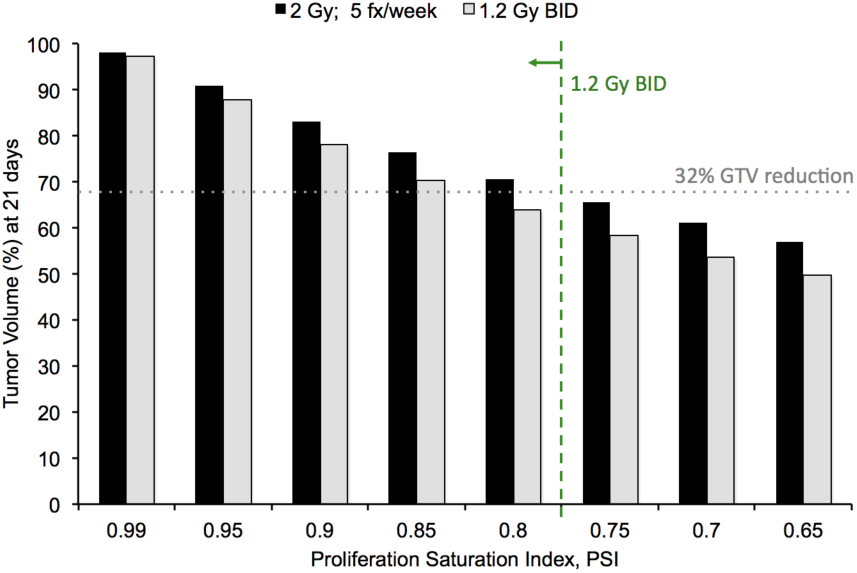
Model of tumor response at 4 weeks by initial pretreatment PSI.

## IV. CONCLUSION

Personalization of radiation fractionation using PSI and mathematical modeling may improve on treatment response and ultimately outcomes in OPC. Validation of the model in additional data sets and ultimately a prospective clinical trial is warranted. PSI can be calculated from delineated volumes on two routinely collected radiological scans (PET/CT at diagnosis and treatment simulation):

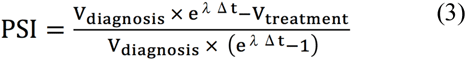

For the analyzed OPC cohort, the PSI-dependent recommended radiation fractionation are as follows:

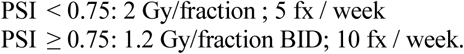

These model predictions are currently evaluated in a prospective clinical trial (NCT03656133).

## Notes

This project was supported in part by a pilot award from the NIH/NCI U54CA193489 “Cancer as a complex adaptive system”.

